# Optimal dynamic empirical therapy in a health care facility: an artificial intelligence approach

**DOI:** 10.1101/603464

**Authors:** Nicolas Houy, Julien Flaig

## Abstract

We propose a solution to the problem of finding an empirical therapy policy in a health care facility that minimizes the cumulative infected patient-days over a given time horizon. We assume that the parameters of the model are known and that when the policy is implemented, all patients receive the same treatment at a given time. We model the emergence and spread of antimicrobial resistance at the population level with the stochastic version of a compartmental model. The model features two drugs and the possibility of double resistance. Our solution method is a variant of the Monte-Carlo tree search algorithm. In our example, this method allows to reduce the cumulative infected patient-days over two years by 22% compared to the best standard therapy.

---

A recent study [14, 49] estimated the burden of antimicrobial resistance in Europe (EU and European Economic Area) in 2015 to be comparable to that of influenza, HIV, and tuberculosis *combined*. Hospitals and other health care facilities were shown to be particularly affected. Of all infections with antibiotic resistant bacteria considered in the study, an estimated 63.5% were associated with health care, accounting for 72.4% of attributable deaths and 74.9% of DALYs (disability adjusted life years).

At the molecular and cellular levels, antimicrobial resistance is mediated by four main mechanisms [23, 50]. An organism may acquire genes allowing it to bypass the metabolic pathways targeted by an antimicrobial agent, to destroy the antimicrobial agent, to alter its target, or to prevent it from reaching its target altogether. At the population level, both within-host and between-host, the spread of resistance is allowed by the selective pressure of antimicrobials [30].

Antimicrobial resistance is a wide-ranging problem, and the relative influence of antimicrobial use in health care facilities on its emergence has sometimes been challenged [22, 47]. Yet several features of health care facility environments make antimicrobial resistance a serious issue in health care facilities [23, 35]. First, patients are highly vulnerable to infections – drug-resistant or not. This may be due to their general health condition (underlying disease, denutrition, previous infections), to breaches into the physical defenses of the body (intravascular devices, tracheal tubes, urinary catheters), or to treatments weakening the immune system. Promiscuity and long or repeated exposure (chronically ill and relapsing patients) increase the risk of infection. Second, and precisely because patients are so vulnerable, antimicrobials are heavily used in health care facilities either as prophylactic treatments or empirically. *Empirical therapies* consist in using combinations of antimicrobials or broad-spectrum antimicrobials at the onset of symptoms, when the infecting organism is not yet known e.g. before test results are available and allow to switch to a more specific treatment. Whether drug-resistance is first acquired by organisms in health care facilities, or mutations first occur in the community and are then imported in health care facilities, exposure to antimicrobials during care (even at residual levels as it has been suggested [19]) selects for drug-resistance with serious consequences for the patients.

Many strategies have been proposed to control health care associated infections, some of which are routine in health care facilities. They include hygiene measures, screening, prophylaxis, isolation of infected patients, and patient and staff cohorting [15]. Since antimicrobials promote the emergence of resistance but are at the same time unavoidable in health care facilities, empirical prescription policies have been proposed to slow and even revert the evolution of resistance [36]. Indeed, antimicrobial resistance is often associated with a fitness cost [1–3, 32] such that a resistant mutant strain’s ability to survive and reproduce in the absence of antimicrobial treatment is reduced compared to that of wildtype drug susceptible strains. Therefore, alternating between antimicrobials or increasing heterogeneity in empirical therapies should reduce the emergence of resistance, at least in theory. Prescription policies based on this idea include *mixing* where antimicrobials are allocated randomly to patients, and *cycling* where antimicrobials are alternated either following a fixed rotation schedule or adaptively.

The merits of cycling and mixing have been largely debated and there is to date no clear-cut empirical evidence that either performs better in general (see [7, 8] and the references therein). As for theoretical studies, it was first shown [9, 27, 45] that mixing performs better than cycling with a fixed period in a vast majority of cases. Later, other authors [40] used a control-theoretic approach to show that a better policy than mixing could be found by relaxing the constraint that drugs should alternate with a fixed period. This approach was criticized on two major accounts [11]. First, that computing an optimal policy depending heavily on the parameters of the problem would require perfect knowledge of such parameters. Second, that a policy without a fixed period would be too complex to implement. That is, put shortly, that trying to compute and implement an optimal policy would be impractical. Given the stakes, the critics advocated a conservative approach to empirical therapy policies. Yet the debate does not seem to be closed as it was shown [8, 41] that the results in [9, 45] could be explained by the parameter values used in these studies (especially symmetries between strains). In addition, the same authors argued that it is impossible to tell *a priori* which of mixing and cycling with a fixed period performs better due to inherent properties of the problem. To be sure, these theoretical arguments do not answer criticisms of optimization made on the ground that it could not be applied in practice, but they clearly speak in favor of exploring this path further – with the provision that an optimization method, however powerful, is ultimately to be appraised in the light of its practical feasibility.

In this article, we propose a solution to the optimization problem that consists in choosing a rotation schedule for, say, new antimicrobials to be used in empirical therapies in a health care facility. For the sake of clarity in the following, we take up a broad definition of empirical therapies including prophylactic treatments. We assume that the parameters of the population and disease dynamics in the health care facility are known. Also, we assume that the rotation schedule is not adaptive, *i.e.* its implementation does not depend on the contingencies of the spread of the disease. Solving the optimization problem in this setting is intended as a first step toward the information rich and personalized strategies pointed to in [8].

This optimization problem is a *dynamic* problem in the sense that the empirical treatment decisions in a given period of time influence the subsequent spread of the disease and emergence of resistance. However, the size and complexity of the problem make common dynamic programming methods ineffective. We circumvent this difficulty by using a variant of the Monte-Carlo tree search algorithm, an algorithm first developed for two-player games in the field of artificial intelligence [12]. Algorithms of the same class have previously found applications in health care, for instance to compute optimal chemotherapy regimens [20, 21].

Our contribution is primarily methodological. Our objective is to illustrate the use of an empirical therapy optimization method. There is no reason to believe *a priori* that alternating between drugs (perhaps following some intricate therapy policy) will yield better results than, say, simple hygiene measures. Yet empirical therapy optimization remains relevant as a part of a global strategy to limit antibiotic resistance. Besides, recent advances in evolutionary biology have opened the way to predicting the emergence of resistance [17], and to exploiting evolutionary interactions between drugs to curb it [7]. On their way to implementation in health care facilities, such novel strategies will certainly have to be optimized at the population level.^1^

The optimization algorithm implemented in this study is applied to a mathematical model of the emergence and spread of drug resistance in a health care facility. Mathematical modeling and numerical simulation are natural tools for infection control in health care facilities. We refer to [18] for an early overview of this topic and to [53] for a review. Flexibility is certainly the main advantage of mathematical modeling as it allows to test and compare control strategies at almost no cost and with little practical or ethical constraints. This is all the more true when designing empirical therapy policies given the large number of available alternatives (see again the arguments made in [8]).^2^ In this line, our method is close to *in silico* clinical trials (see [33] for instance).

While antimicrobial resistance is but one issue among others in health care facility infection control, its complexity received special attention from modelers. Reviews and categorizations of models of emergence and spread of antimicrobial resistance can be found in [44, 48]. In this study, we build upon models of antimicrobial resistance that were developed to investigate empirical therapy policies. These models are compartmental models at the between-host level. They do not give an explicit account of within-host dynamics. Early studies in this stream of literature include [4, 5, 26, 31] with models featuring a single drug. The discussion of the relative merits of mixing and cycling found in [9, 45] and [40] relies on models derived from [10, 28, 29]. Consequently, we use a variant of the same models. Similar variants have previously been used to study adaptive and adjustable strategies [24, 55], to investigate the role of hospital size in the spread of infections [25], to understand the implications at the population level of resistance acquisition mechanisms reported *in vitro* [37], to give justification for the widely used combination therapies [13], to discuss potential adverse effects of antibiotic restrictions [38], and to compare empirical therapy policies over ranges of population dynamics parameters [51, 52].

As we are investigating a nonlinear phenomenon in a small population (such as a hospital unit), we will use stochastic simulations of the model. We refer to [16] for a striking theoretical illustration of stochastic effects in a hospital setting, and to [24, 25] for practical implications. Specially in the context of optimization, stochastic effects cannot be overlooked.

We expose the materials and methods used in the study in Section 1. The model of emergence and spread of resistance in a health care facility is presented in Section 1.1. The optimization algorithm is introduced in Section 1.2. We show our results in Section 2: first, the performance of the optimal policy computed with our optimization algorithm (Section 2.1), then the performance of a close policy with the advantage of a fixed switching period (Section 2.2). Section 3 concludes the study.

## 1. Materials and methods

### 1.1 Model

We describe with an epidemiological compartmental model the emergence and spread of a microbial disease and of resistant strains in a small health care facility or a single hospital ward. Two drugs are available, drug 1 and drug 2. At any point in time, patients are either infected with a wild-type strain susceptible to both drugs, infected with a strain resistant to drug 1 and susceptible to drug 2 (1-resistant), infected with a strain resistant to drug 2 and susceptible to drug 1 (2-resistant), infected with a strain resistant to both drugs (12-resistant), or uninfected with any of the above. We assume that the different strains do not coexist in a single host.

As was noted elsewhere [56], a new strain will take over a resident population of organisms following two distinct processes. First, the newcomer must get a foothold in the host. In our model, a new strain appears in a host either through *de novo* mutation, or through migration, that is contagion. Horizontal gene transfers, another widely studied mode of emergence of new strains [6, 54], are overlooked in this study. Once it has appeared within a host, the new strain must compete with the resident strains to survive and reproduce. When the patient is not infected by one of the four pathogenic strains, the new pathogenic strain must compete with the commensal microflora. We assume that commensal organisms cannot be transmitted, and that they are not impacted by antimicrobial treatments.^3^ As a result of the two processes, an equilibrium is finally reached where one strain takes over the other.

Drug resistant organisms may incur a fitness cost in each of the two processes [3]. Accordingly, we include two distinct fitness costs in our model. *c*_*i*_*∈* [0, 1], *i* ∈ {1, 2, 12}, denotes the fitness cost on transmission of the *i*-resistant strain. The transmission rate of a *i*-resistant strain is weighed by 1 *- c*_*i*_. We normalize the fitness cost *c*_*S*_ of the wild-type susceptible strain to 0. *s*_*i*_ ∈ [0, 1], *i* ∈ {1, 2, 12}, denotes the fitness cost on competitive performance of the *i*-resistant strain in the absence of adequate treatment. Again, we normalize the fitness cost *s*_*S*_ of the wild-type susceptible strain in the absence of treatment to 0. Also, the fitness cost *s*_*X*_ of the commensal microflora is normalized to 1. We assume that the fitness cost on competitive performance of a strain that receives adequate treatment is equal to 1. In the literature, the fitness cost *s*_*i*_ is often interpreted as a selection coefficient, that is a reduction in growth rate due to resistance, so that in most cases only new alleles providing a strict advantage over the resident population have a non-zero probability of fixation (takeover). In our illustrative framework, we allow the fixation of neutral and deleterious mutations. However, we assume that a strain cannot colonize a treated patient unless it is resistant to the treatment. If the patient is not treated or if the new strain is resistant to the treatment, the rate of takeover of the resident strain by the new strain is given by

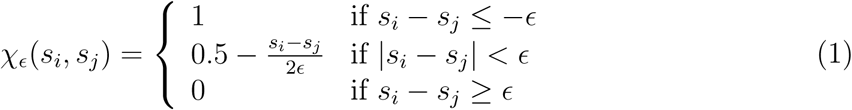

where *s*_*i*_ is the fitness cost on competitive performance of the new strain, and *s*_*j*_ the fitness cost on competitive performance of the resident strain. In the limit ϵ *→* 0, a new strain takes over the population if and only if it has a competitive advantage over the resident population.^4^ Function *χ*_*ϵ*_ allows us to model the relative competitive performance of strains in a flexible and consistent way, while avoiding to model within-host dynamics explicitly. A useful graphical summary of the evolutionary dynamics of resistance in the presence and absence of treatment can be found in [56]. We refer the interested reader to [39] for a brief historical overview of the derivation of fixation probabilities. As a final remark about fitness costs and selection, we point out that our framework allows to model fitness costs of drug *susceptibility* as have been reported in several organisms [46], even if we made the more common assumption of fitness costs associated with drug *resistance*.

At each time, patients can receive a treatment. Three treatments are available to the clinician: drug 1 in monotherapy (treatment 1), drug 2 in monotherapy (treatment 2), and a combination of drugs 1 and 2 (treatment 12). Patients can also be left without a treatment (treatment 0). Decision variable *f*_*i*_, *i* ∈ {0, 1, 2, 12}, is equal to 1 when patients receive treatment *i* and to 0 otherwise. Notice that at each time exactly one of the *f*_*i*_’s, *i* ∈ {0, 1, 2, 12}, is equal to 1 and the others are equal to 0.

Our compartmental model is illustrated in Figure 1. The events corresponding to moves from and to compartments are summarized in Table 1, and the parameters are shown in Table 2 along with their default values.

**Table 1:** Events considered in the model.

| Population changes | Rate |
| --- | --- |
| *Admissions and discharges* |  |
| $S \leftarrow S + 1$ | $N\mu m_S$ |
| $R_i \leftarrow R_i + 1, i \in \{1, 2, 12\}$ | $N\mu m_i$ |
| $X \leftarrow X + 1$ | $N\mu \left(1 - \sum_{i \in \{S, 1, 2, 12\}} m_i\right)$ |
| $S \leftarrow S - 1$ | $\mu S$ |
| $R_i \leftarrow R_i - 1, i \in \{1, 2, 12\}$ | $\mu R_i$ |
| $X \leftarrow X - 1$ | $\mu X$ |
| *De novo mutations* |  |
| $S \leftarrow S - 1, R_i \leftarrow R_i + 1, i \in \{1, 2\}$ | $\nu_i [f_0 \chi_\epsilon(s_i, s_S) + f_i \chi_\epsilon(s_i, 1)] S$ |
| $S \leftarrow S - 1, R_{12} \leftarrow R_{12} + 1$ | $\nu_{12} [f_0 \chi_\epsilon(s_{12}, s_S) + (f_1 + f_2 + f_{12}) \chi_\epsilon(s_{12}, 1)] S$ |
| $R_i \leftarrow R_i - 1, R_{12} \leftarrow R_{12} + 1, i \in \{1, 2\}$ | $\nu_{(3-i)} [(f_0 + f_i) \chi_\epsilon(s_{12}, s_i) + (f_{(3-i)} + f_{12}) \chi_\epsilon(s_{12}, 1)] R_i$ |
| *Infections and recoveries* |  |
| $X \leftarrow X - 1, S \leftarrow S + 1$ | $f_0 \beta (1 - c_S) \chi_\epsilon(s_S, 1) X S / N$ |
| $X \leftarrow X - 1, R_i \leftarrow R_i + 1, i \in \{1, 2\}$ | $(f_0 + f_{(3-i)}) \beta (1 - c_i) \chi_\epsilon(s_i, 1) X R_i / N$ |
| $X \leftarrow X - 1, R_{12} \leftarrow R_{12} + 1$ | $\beta (1 - c_{12}) \chi_\epsilon(s_{12}, 1) X R_{12} / N$ |
| $S \leftarrow S - 1, X \leftarrow X + 1$ | $[\tau(f_1 + f_2 + f_{12}) + \gamma] S$ |
| $R_i \leftarrow R_i - 1, X \leftarrow X + 1, i \in \{1, 2\}$ | $[\tau(f_{(3-i)} + f_{12}) + \gamma] R_i$ |
| $R_{12} \leftarrow R_{12} - 1, X \leftarrow X + 1$ | $\gamma R_{12}$ |
| *Superinfections* |  |
| $S \leftarrow S - 1, R_i \leftarrow R_i + 1, i \in \{1, 2\}$ | $\sigma \beta (1 - c_i) [f_0 \chi_\epsilon(s_i, s_S) + f_i \chi_\epsilon(s_i, 1)] S R_i / N$ |
| $R_i \leftarrow R_i - 1, S \leftarrow S + 1, i \in \{1, 2\}$ | $\sigma \beta (1 - c_S) f_0 \chi_\epsilon(s_S, s_i) S R_i / N$ |
| $S \leftarrow S - 1, R_{12} \leftarrow R_{12} + 1$ | $\sigma \beta (1 - c_{12}) [f_0 \chi_\epsilon(s_{12}, s_S) + (f_1 + f_2 + f_{12}) \chi_\epsilon(s_{12}, 1)] S R_{12} / N$ |
| $R_{12} \leftarrow R_{12} - 1, S \leftarrow S + 1$ | $\sigma \beta (1 - c_S) f_0 \chi_\epsilon(s_S, s_{12}) S R_{12} / N$ |
| $R_i \leftarrow R_i - 1, R_{12} \leftarrow R_{12} + 1, i \in \{1, 2\}$ | $\sigma \beta (1 - c_{12}) [(f_0 + f_i) \chi_\epsilon(s_{12}, s_i) + (f_{(3-i)} + f_{12}) \chi_\epsilon(s_{12}, 1)] R_i R_{12} / N$ |
| $R_{12} \leftarrow R_{12} - 1, R_i \leftarrow R_i + 1, i \in \{1, 2\}$ | $\sigma \beta (1 - c_i) (f_0 + f_i) \chi_\epsilon(s_i, s_{12}) R_i R_{12} / N$ |
| $R_i \leftarrow R_i - 1,$<br>$R_{(3-i)} \leftarrow R_{(3-i)} + 1, i \in \{1, 2\}$ | $\sigma \beta (1 - c_{(3-i)}) [f_0 \chi_\epsilon(s_{(3-i)}, s_i) + f_{(3-i)} \chi_\epsilon(s_{(3-i)}, 1)] R_i R_{(3-i)} / N$ |

**Table 2:** Model parameters.

| Parameter | Unit | Description | Value |
| --- | --- | --- | --- |
| *Admissions and discharges* |  |  |  |
| $N$ | – | Average number of patients. | 40 |
| $\mu$ | day<sup>-1</sup> | Turnover rate. | 0.1 |
| $m_S$ | – | Proportion of incoming patients infected with the susceptible strain. | 0.1 |
| $m_1$ | – | Proportion of incoming patients infected with the strain resistant to drug 1. | 0.05 |
| $m_2$ | – | Proportion of incoming patients infected with the strain resistant to drug 2. | 0.01 |
| $m_{12}$ | – | Proportion of incoming patients infected with the strain resistant to drugs 1 and 2. | 0.005 |
| *De novo mutations and fitness costs* |  |  |  |
| $\nu_i$ | day<sup>-1</sup> | Rate of acquisition of resistance to treatment $i$ through *de novo* mutation. $i \in \{1, 2, 12\}$ | 0.005 |
| $c_i$ | – | Cost on infectivity of resistance to treatment $i$, $i \in \{1, 2, 12\}$. | 0.2 |
| $c_S$ | – | Cost on infectivity of susceptible organisms. | 0 |
| $s_i$ | – | Cost on competitive performance of resistance to treatment $i$, $i \in \{1, 2, 12\}$. | 0.2 |
| $s_S$ | – | Cost on competitive performance of susceptible organisms. | 0 |
| $s_X$ | – | Cost on competitive performance of commensal organisms. | 1 |
| $\epsilon$ | – | Slope parameter in function $\chi_\epsilon$. | 0.5 |
| *Infections, superinfections, and recoveries* |  |  |  |
| $\beta$ | day<sup>-1</sup> | Transmission rate. | 0.5 |
| $\sigma$ | – | Relative rate of superinfection. | 1 |
| $\gamma$ | day<sup>-1</sup> | Rate of spontaneous recovery. | 0.1 |
| $\tau$ | day<sup>-1</sup> | Rate of recovery under appropriate treatment. | 0.3 |

**Figure 1:**
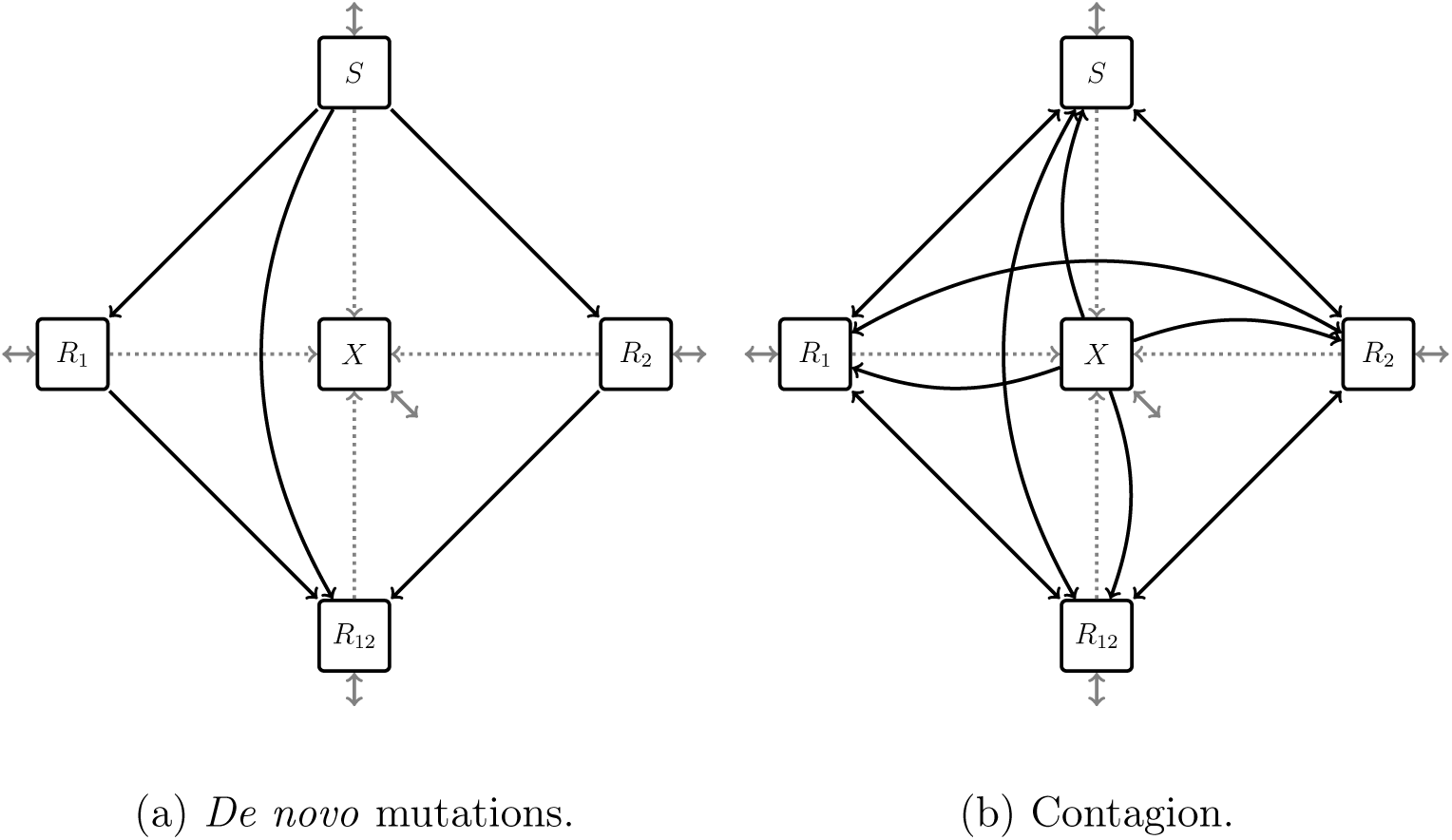
For clarity we display *de novo* mutations *(black, panel (a))* and contagion *(black, panel (b))* on two separate graphs. *Gray:* admissions and discharges. *Dotted gray:* recovery.

*S, R*_*i*_ and *X* denote the number of patients infected with the susceptible strain, the *i*-resistant strains (*i* ∈ {1, 2, 12}), and the number of uninfected patients respectively. It is assumed that infection with one of the four pathogenic strains considered in the model is not the primary cause of admission, so the admission and discharge rates are independent of health status. When patients are infected at the time of admission, the symptoms appear following the start of care. This is characteristic of endogenous infections, where a previously harmless organism is introduced by health interventions in body parts where it causes damage. For simplicity, we assume that the spread of infections in the community outside the health care facility is exogenous to the problem – the proportion of incoming patients infected with each strain is set as a parameter *m*_*i*_, *i* ∈ {*S*, 1, 2, 12}.^5^ Also, we assume that infected patients do not incur additional mortality or morbidity.

Infected individuals recover spontaneously at rate *γ*, or at rate *τ* if they receive adequate treatment. Rates *γ* and *τ* do not depend on the infecting strain. When a treatment is adequate, the rate *τ* does not depend on the antimicrobial or combination of antimicrobials (there are no drug interactions).

*De novo* mutations conferring drug resistance (Figure 1a) occur on two different genes and at constant rates, as illustrated in Figure 2. We assume that a mutation conferring 1-resistance (resp. 2-resistance) cannot take over the pathogenic population in a host receiving treatment 2 or 12 (resp. treatment 1 or 12), unless the resident population (and so the mutant) is already 2-resistant (resp. 1-resistant). The level of selective pressure depends on whether the resident population receives adequate treatment or not, as modeled by fitness c osts a nd r ate f unction *χ* _*ϵ*_. S ince t here i s n o c oexistence o f p athogenic strains within a host, acquisition of 1-resistance (resp. 2-resistance) by an organism infecting a host in *R*_2_ (resp. *R*_1_) leads necessarily to 12-resistance. Finally, our model assumes no back-mutations, no compensatory mutations, and no co-selection.

**Figure 2:**
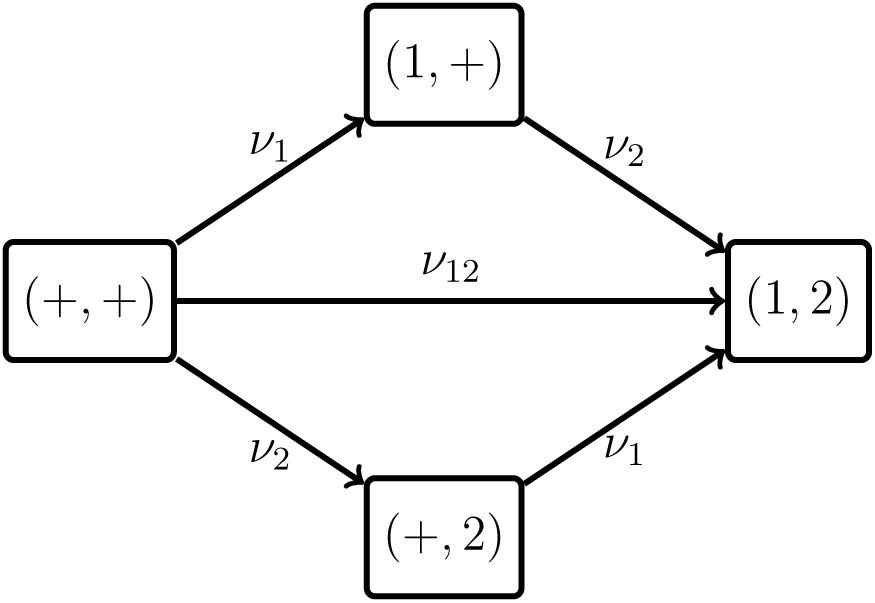
The four genotypes included in our model with mutation rates. The arrows represent transitions between genotypes. The two considered genes are separated by a comma. ‘+’ represents a wild-type allele. ‘1’ and ‘2’ represent alleles giving resistance to drugs 1 and 2 respectively. (+, +): wild-type susceptible. (1, +): 1-resistance only. (+, 2): 2-resistance only. (1, 2): resistance to both drugs. Adapted from [55].

Infection and superinfection (Figure 1b) are the second mode of acquisition of a new strain in our model. Notice that because all patients receive identical treatment, a strain under adequate treatment cannot be transmitted before it is cleared.

A patient already infected with a pathogenic strain may become superinfected with a new strain. Treatments offer protection so a strain cannot be transmitted to a patient receiving a treatment, unless the strain is resistant to this treatment. If it is, and just like in the case of *de novo* mutations, its success at taking over the resident infecting population is represented by rate function *χ*_*ϵ*_. This success depends on the treatment currently received by the host, and on the resident strain ability to fight off i nvasion g iven t hat treatment. In this instance of the model, we make the assumption that the transmission to an already infected host when the competitive advantage over the resident strain is high (*χ*_*ϵ*_ = 1) is equal to the transmission rate to an uninfected individual. Put differently, the relative rate of superinfection *σ* is set to 1. This needs not always be the case (e.g. because of the patient’s immune response to the first infection) and we give a general formulation including *σ* in Table 1 for later reference.

### 1.2 Optimization algorithm

In our scenario, we must decide on an empirical therapy policy knowing only the population dynamics parameters of the disease and of the health care facility. In particular, we do not allow for strategies that are contingent on observations of the actual evolution of the disease. Decision heuristics based on the evolution of the number of patients infected with each strain for instance have shown promising results in theoretical studies [8, 24], but this information is not always readily available to clinicians, e.g. when a rotation schedule is to be chosen in advance for new antimicrobials that have not been used before. We leave the important issue of optimization when more information is available to future studies. Let us now turn to a more detailed explanation of our algorithm.

We compute an empirical therapy policy by arranging the use of the four treatment described in the previous section over an horizon *H* of two years (720 days). In the following, a *policy* denotes a sequence of treatments over a period of time that needs to be specified (administering a single treatment over the period is a valid policy). Our objective is to minimize the cumulative infected patient-days over *H*. We call OPTI the policy computed with our optimization algorithm. In OPTI, the treatment can be switched every Δ*h* = 30 days. Therefore, *L* = 24 *decisions* are to be made. Available decisions for each period of length Δ*h* consist in administering either of the following treatments during each period: treatment 1, treatment 2, treatment 12, or no treatment. We define the corresponding standard policies: MONO-1 which consists in always administering treatment 1 over the specified period of time, MONO-2 which consists in always administering treatment 2, COMBO which consists in always administering treatment 12, and NONE which consists in never administering any drug. Of course, the merit of the immediate decision for one period depends on the subsequent spread of the infection, that is on the subsequent decisions that are yet unknown at the moment of decision. In the algorithm, decisions in each period are made by assuming that a *default policy* will be applied subsequently over *H -*Δ*h* days. Notice that this horizon is rolling so that there are no border effects. Decisions are evaluated on the basis of the cumulative infected patient-days over the rolling horizon of length *H* days (the decision during Δ*h* days followed by the default policy during *H -* Δ*h* days). To evaluate decisions while taking stochastic effects into account, we run *n*_*S*_ = 2000 stochastic simulations of the model for each possible decision over the rolling horizon and choose the policy that minimizes the average cumulative infected patient-days.

The performance of the algorithm depends on the fine-tuning of a number of parameters. The length Δ*h* of a period can be adjusted. If it is too short (e.g. one day), stochastic effects come to dominate the effect of the decision and the optimization fails. Keep also in mind that switching treatments too often can be impractical in a health care facility. The rolling horizon does not need to have the same length as the policy we are computing. It can be shortened, for instance if computing resources are limited, but at the expense of the performance of the computed policy. *n*_*s*_ must be chosen so that the algorithm is not too sensitive to stochastic effects (*n*_*s*_ too small), but still does not require unreasonably high computing power (*n*_*s*_ too large). Also, different default policies will yield different solutions (“optimal” policies). Therefore several default policies need to be experimented and improved upon. We tested different values for the parameters of our algorithm, and chose those yielding the best results.

Our optimization method is presented more formally in Algorithms 1–3. Algorithm 1 shows the procedure stochSimu updating a stochastic simulation of the model over a time horizon given a treatment policy. Function evaluateDecision computes the average cumulative infected patient-days over *n*_*S*_ simulations obtained by applying a sequence of two policies over two subsequent time horizons. Algorithm 2 is the core of the method. It allows to compute a solution policy given a default policy as explained above. Algorithm 3 shows how the method is initialized, and how different default policies are tried in order to improve the solution. We initialize all simulations by running the model over a 30-year period under the NONE policy (no treatment). We first use NONE, MONO-1, MONO-2, and COMBO as default policies. Thus, we obtain four solution policies. The best solution policy out of the four is then used as default policy to compute a new solution. We iterate by using the last solution as default policy to compute the next solution until the quality of the solution deteriorates.

## 2. Results

### 2.1 Performance of our optimal policy OPTI

Figure 3 illustrates the optimal policy OPTI computed with our algorithm. Treatment 1 is never used, and the policy consists mostly in switching between treatment 0 (no treatment) and treatment 12 (both drugs).

**Figure 3:**
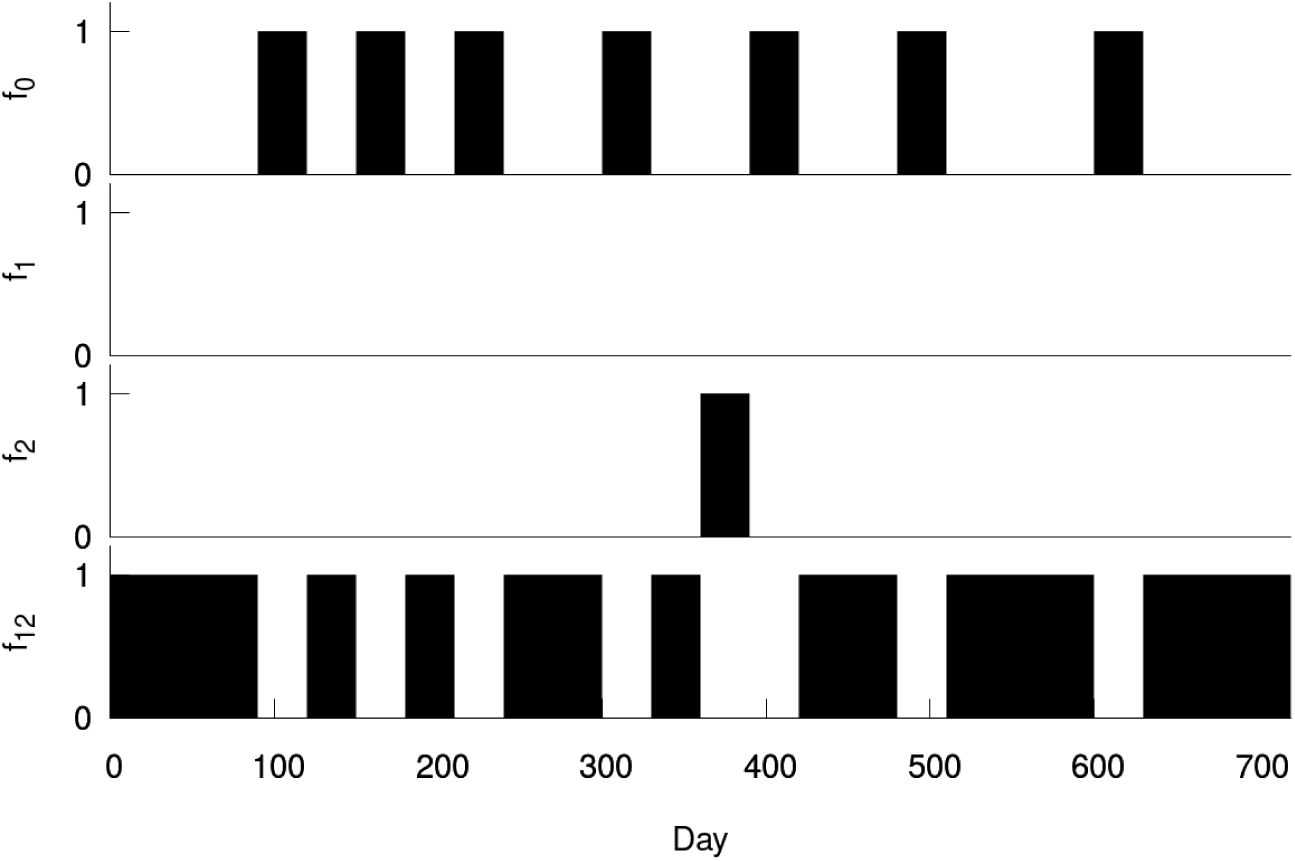
Optimal policy OPTI.

Once OPTI has been designed, we run 400 stochastic simulations of the model under this policy. We also run 400 simulations under policies NONE, COMBO, MONO-1, and MONO-2 defined in Section 1.2. In Figure 4, we compare the performance the different policies over two years (720 days). We show the performance of cycling policies in order to compare our results with those of standard policies found in the literature. The cycling policy CYC-*n* consists in alternating *n* days of treatment 1 with *n* days of treatment 2. Without treatment (NONE), we obtain an average of 18,215.4 (95% CI: 18,126.2 – 18,304.6) cumulative infected patient-days over two years. The combination therapy COMBO performs better than cycling policies with an average of 12,314.5 (95% CI: 12,122.3 – 12,506.8) cumulative infected patient-days over two years, hence allowing a 32.4% reduction compared to the NONE policy. The best cycling policy is CYC-1 with a 31.5% reduction in the average cumulative infected patient-days compared to NONE (12,462.1, 95% CI: 12,266.6 – 12,657.7). OPTI almost halves the average cumulative infected patient-days obtained under no treatment with a 47% decrease: average of 9,629.61 (95% CI: 9,462.96 – 9,796.25) cumulative infected patient days. Compared to COMBO, OPTI reduces the average cumulative infected patient-days over two years by 22%.

**Figure 4:**
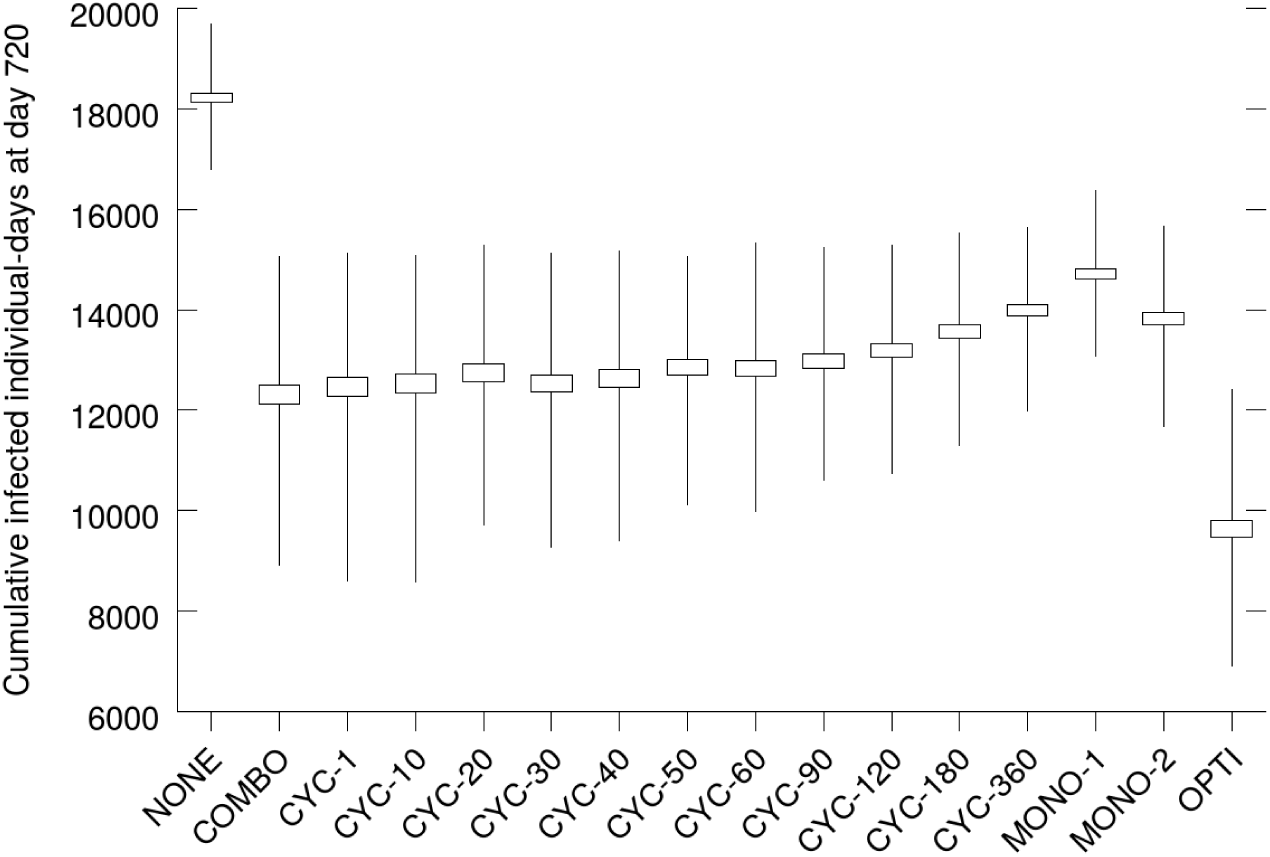
Box plot of the cumulative infected patient-days (400 simulations). *Box:* 95% confidence interval. *Whiskers:* 5^th^ and 95^th^ percentiles of the simulations.

**Figure 5:**
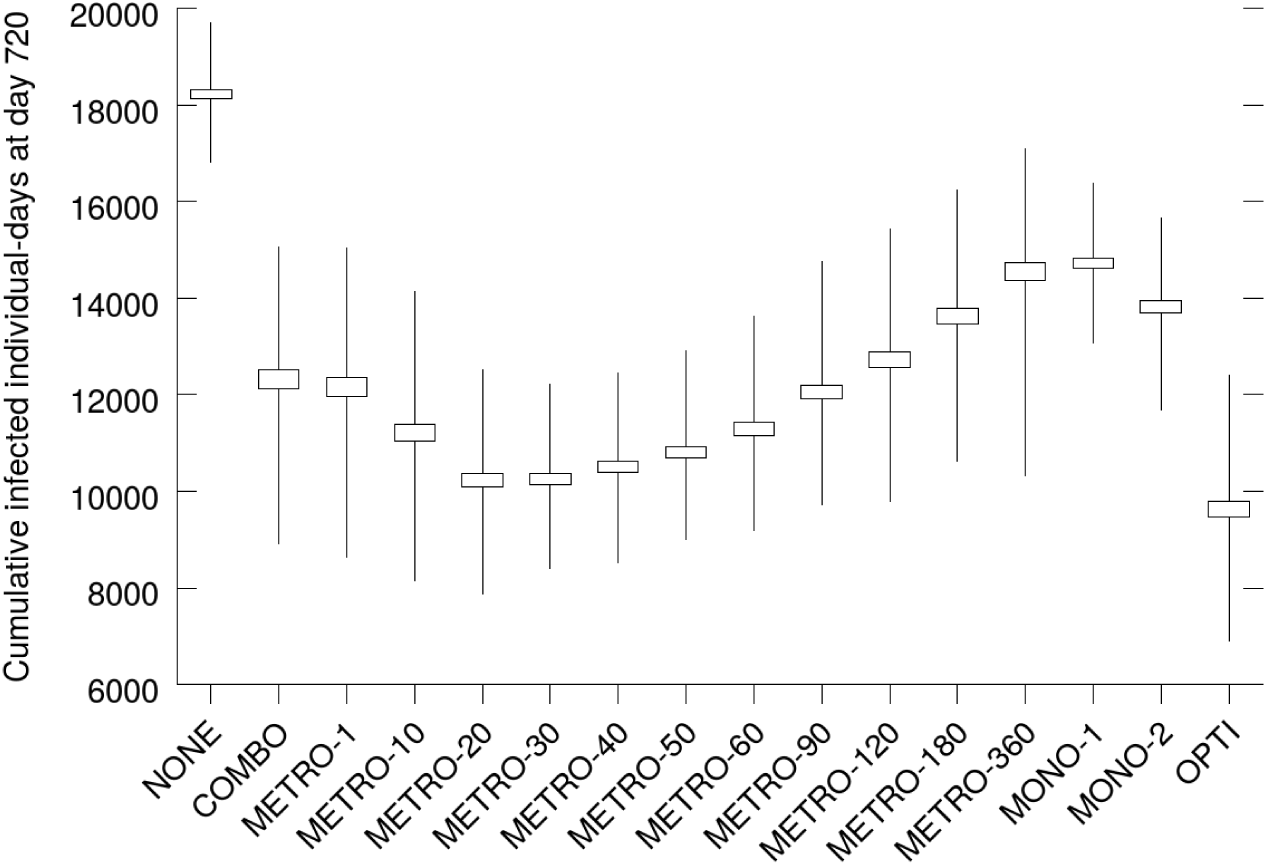
Box plot of the cumulative infected patient-days (400 simulations). *Box:* 95% confidence interval. *Whiskers:* 5^th^ and 95^th^ percentiles of the simulations.

#### Algorithm 1 Simulation of the model, and policy and decision evaluation

~~~
1: **procedure** stochSimu<(*s, π, H, h*)
        ▹ Update simulation *s* by running the stochastic model with treatment policy *π* starting at date *h* and over horizon *H*.
2: **simulate** *s* starting from its current state, using the model presented in Table 1 with the *f*_*i*_’s, *i* ∈ {0, 1, 2, 12}, varying according to *π* starting at date *h* and over *H*
3: **end procedure**
4: **function** EVALUATEDECISION(*S, π*_1_, *H*_1_, *π*_2_, *H*_2_)
        ▹ Evaluate the performance over simulation set *S* of using treatment policy *π*_1_ over horizon *H*_1_ (the immediate decision) followed by policy *π*_2_ starting at date *H*_1_ and over *H*_2_ (the default policy), and return a score.
5:   **for all** *s* ∈ *S* **do**
6:      STOCHSIMU(*s, π*_1_, *H*_1_, 0)
7:      STOCHSIMU(*s, π*_2_, *H*_2_, *H*_1_)
8:   **end for**
9:   **return** average cumulative infected patient-days over *S*
10: **end function**
~~~

#### Algorithm 2 Policy computation given a default policy

~~~
1: **function** COMPUTEPOLICY(*S*, Π, *π*_*default*_, Δ*h, L*)
        ▹ Compute a treatment policy over horizon *H* = Δ*h × L* by choosing the best policy in set Π for each interval Δ*h* assuming that default policy *π*_*default*_ is used next over horizon *H* - Δ*h*.
2:   *H ←* Δ*h* × *L*
3:   *policy ←* empty vector of size *L*
4:   **for** *l ←* 0 to *L* - 1 by 1 **do**
5:      **for all** *π* ∈ Π **do**
6:        *S*_*π*_ *←* a copy of *S*
7:        *score*_*π*_ EVALUATEDECISION(*S*_*π*_, *π*, Δ*h, π*_*default*_, *H -* Δ*h*)
8:      **end for**
9:      *π*^*\**^ ← argmax_*π*∈Π_ *score*_*π*_
10:     *policy*[*l*] *π←*^*\**^
11:     **for all** *s* ∈ *S* **do** ▹ Update all simulations in *S*.
12:       STOCHSIMU(*s, π*^*\**^, Δ*h*)
13:     **end for**
14:   **end for**
15:   **return** *policy*
16: **end function**
~~~

#### Algorithm 3 Optimization protocol

~~~
**Require:** a stochastic model as presented in Table 1, a horizon *H* split into *L* segments of size Δ*h*.
1: Π ← {NONE, MONO-1, MONO-2, COMBO} ▹ Set of available decisions for each segment of size Δ*h*.
2: Π_*default*_ ← {NONE, MONO-1, MONO-2, COMBO} ▹ Initial set of default policies to be assumed heuristically.
3: *S*_*ini*_ ← a set of *n*_*S*_ simulations (initial conditions at this point)
4: **for all** *s* ∈*S*_*ini*_ **do**                    ▹ Initialization.
5:      STOCHSIMU(*s*, NONE, 30 years)
6: **end for**
7: **for all** *π*_*default*_ ∈ Π_*default*_ **do**               ▹ Try all default policies in Π_*default*_.
8:      *S ←* a copy of *S*_*ini*_
9:      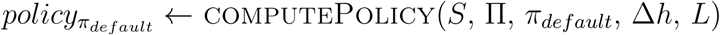
10:     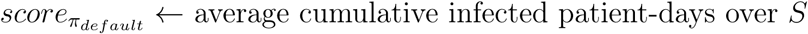
11: **end for**
12: 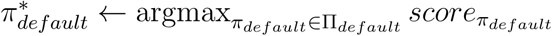
13: 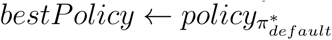
14: 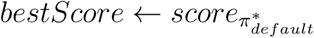
15: *done ← FALSE*                        ▹ Try to improve on *bestPolicy*.
16: **while** !*done* **do**
17:     *S ←* a copy of *S*_*ini*_
18:     *policy ←* COMPUTEPOLICY(*S*, Π, *bestPolicy*, Δ*h, L*)                      ▹ *bestPolicy* is used as default policy.
19:     *score ←* average cumulative infected patient-days over *S*
20:     **if** *score > bestScore* **then**
21:        *bestScore ← score*
22:        *bestPolicy ← policy*
23:     **else**
24:        *done ← TRUE*
25:     **end if**
26: **end while**
27: OPTI *← bestPolicy*
~~~

In Figure 6 we display the evolution in time of the cumulative infected patient-days for OPTI and COMBO. Both policies show similar performance until approximately day 250. We surmise that by day 250, the influence of initial conditions has vanished so the remaining of the curves reflects the divergence between a population treated with OPTI and a steady state population treated with COMBO. We show the evolution of the population in each compartment under policies NONE, COMBO, and OPTI in Figures A.1–A.2 in Appendix.

**Figure 6:**
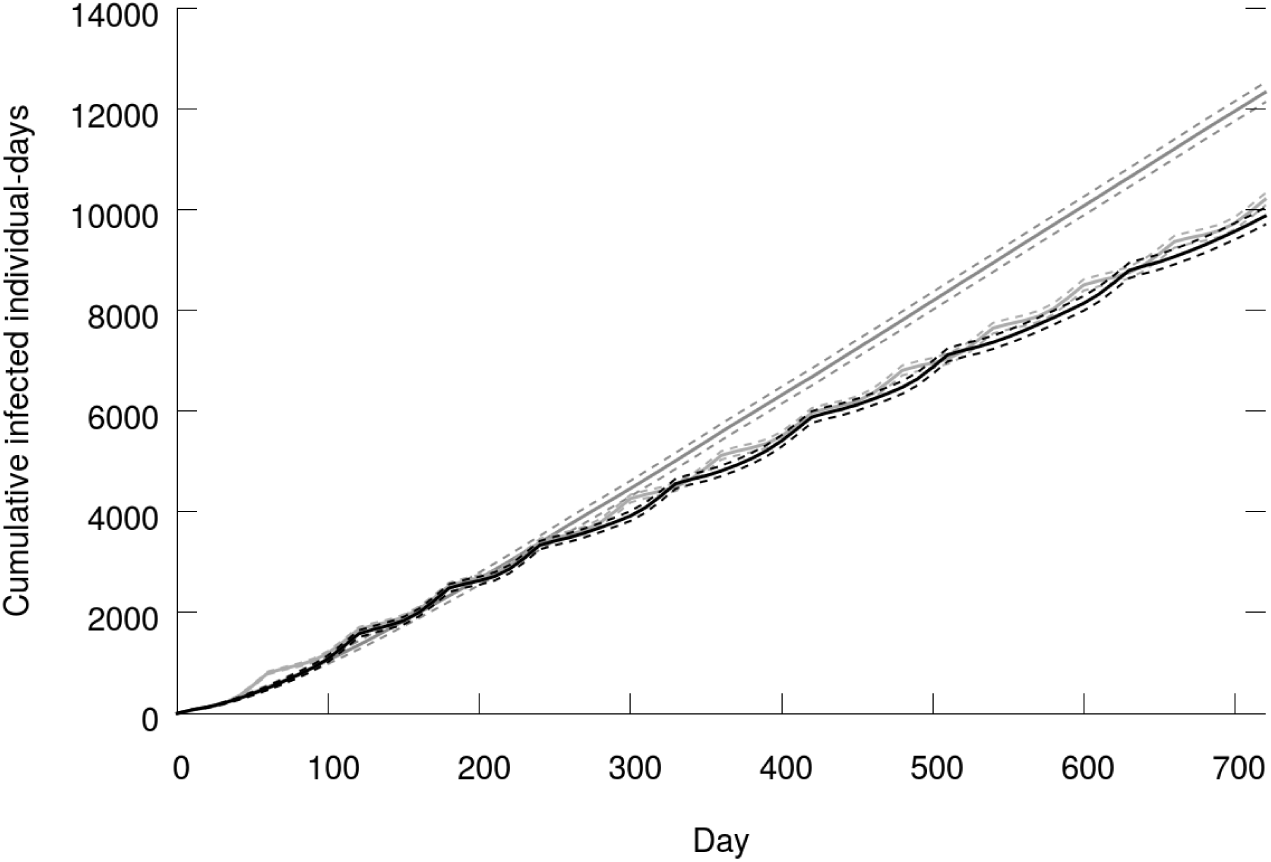
Average cumulative infected patient-days as a function of time (400 simulations). *Black:* OPTI. *Gray (above):* COMBO. *Gray (below):* METRO-30. *Dashed:* 95% CI.

### 2.2 Metronomic therapy policies as alternatives to OPTI

A major drawback of a policy such as OPTI is its irregularity making its use inconvenient in a health care facility. We saw from Figure 3 that OPTI is close to alternating between treatment 12 and treatment 0. Let us compare OPTI with policies that are close to it in this respect, but with the advantage of regularity. We define metronomic policies that consist in switching between treatment 12 and treatment 0 with a fixed period. In METRO-*n*, the switch occurs every *n* days.

The outcomes of METRO-*n* policies are displayed in Figure 6. Among metronomic policies, METRO-20 shows the best result with an average of 10,233.7 (95% CI: 10,090.2 – 10,377.1) cumulative infected patient-days over two years. It is yet arguable that METRO- 30 would be more convenient to implement (one switch every month). Because the average cumulative infected patient-days is only 0.2% (non significantly at the 95% threshold) higher with METRO-30 (10,255, 95% CI: 10,135.7 – 10,374.3) and because METRO-30 shows the advantage of having the same switching dates as OPTI, we will retain METRO- 30 for the following discussion. The evolution of the population in each compartment under METRO-30 as a function of time is given in Figure A.4 in appendix.

OPTI decreases the cumulative infected patient-days by 6% (significantly at the 95% threshold) compared to METRO-30. The close performance of OPTI and METRO-30 is further illustrated in Figure 6. Notice that even though they are always close, there is no time horizon for which METRO-30 dominates OPTI. The close performance of OPTI and METRO-30 has two consequences. First, it shows that even if policies obtained with optimization methods can be impractical, they may still lead the way to more convenient solutions that were first not considered. This might prove especially useful in more com-plex settings, say with more than two drugs and drug interactions or specific constraints. Second, metronomic policies seem to deserve closer investigations. To our knowledge, investigations of periodic empirical therapy policies have mostly been focused on cycling policies that alternate between drugs. Clearly, not treating infected patients may imply some ethical discussions. However, when this option is on the table, metronomic empirical therapy policies should be given proper attention.

## 3 Conclusion

We tackled the problem of designing an empirical therapy policy at the population level in a health care facility. An empirical therapy policy specifies which antimicrobial or combination of antimicrobials should be used empirically at each time. The problem is to choose a policy allowing to cure patients while at the same time avoiding the emergence of antimicrobial resistant strains. To solve the problem, we built upon existing compartmental models developed over the last 20 years and which describe the emergence and spread of antimicrobial resistance in a health care facility. Our version of the model features two drugs and the possibility of double resistance. We use stochastic simulations of the model.

Most previous studies investigating empirical therapy policies compared the performance of policies over ranges of population dynamics parameters. Researchers strove for empirical therapy policies showing good results in general, that is for the most plausible parameter values. While giving priority to such general solutions might be appropriate in some cases, the approach has shown limits as it did not yield unambiguous results as to whether a policy dominates the others in all relevant cases.

In this study, we looked at the problem from another perspective by assuming that the population dynamics parameters are known to the decision maker. Given these parameters, a policy that minimizes the cumulative infected patient-days must be designed. We borrowed our solution method from the field of artificial intelligence. In our setting, the policy computed with the algorithm allows a 22% reduction in the average cumulative infected patient-days over two year compared to the widespread combination therapy. From this policy, considering regularity constraints, we could derive a policy with a fixed period and a close performance. This new policy that we called metronomic has never been suggested in the literature to the best of our knowledge.

The results shown here are only illustrative of the performance of our method. This performance will depend on the problem in hand, and specially on the population and disease dynamics parameters. Yet, we strongly believe that our approach, which is close to *in silico* clinical trials, and the purposely highly flexible solution method presented in this study will contribute to determine implementable optimal empirical therapy policies in more complex and realistic settings.

## A Additional figures

**Figure A.1:**
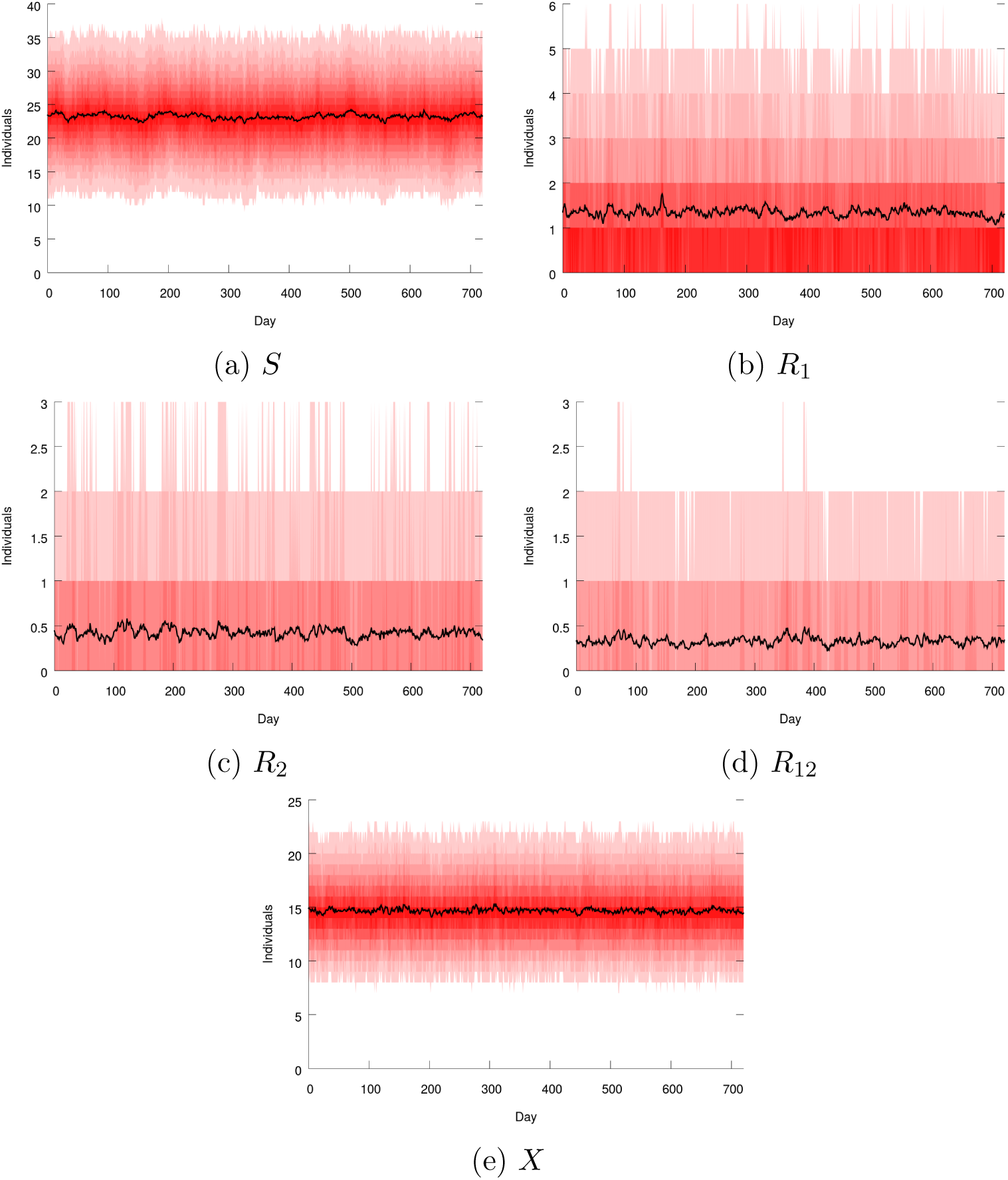
Number of patients in each compartment as a function of time when no treatment is used (NONE policy). *Black:* average over 400 simulations. *Shades of red: i*^th^ and (100 *- i*)^th^ percentiles for *i* from 5 to 50 by increments of 5.

**Figure A.2:**
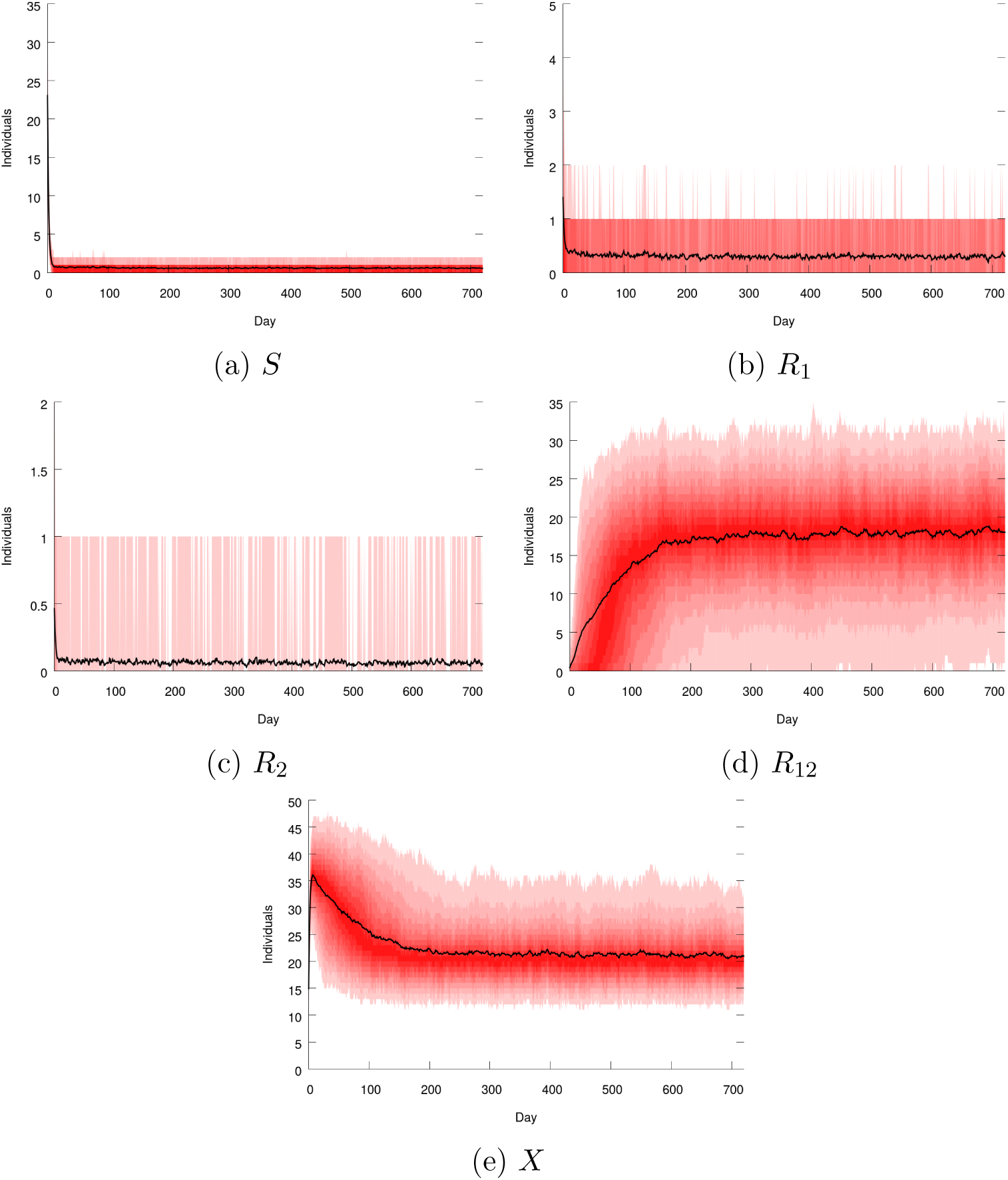
Number of patients in each compartment as a function of time when the COMBO policy is used. *Black:* average over 400 simulations. *Shades of red: i*^th^ and (100 *- i*)^th^ percentiles for *i* from 5 to 50 by increments of 5.

**Figure A.3:**
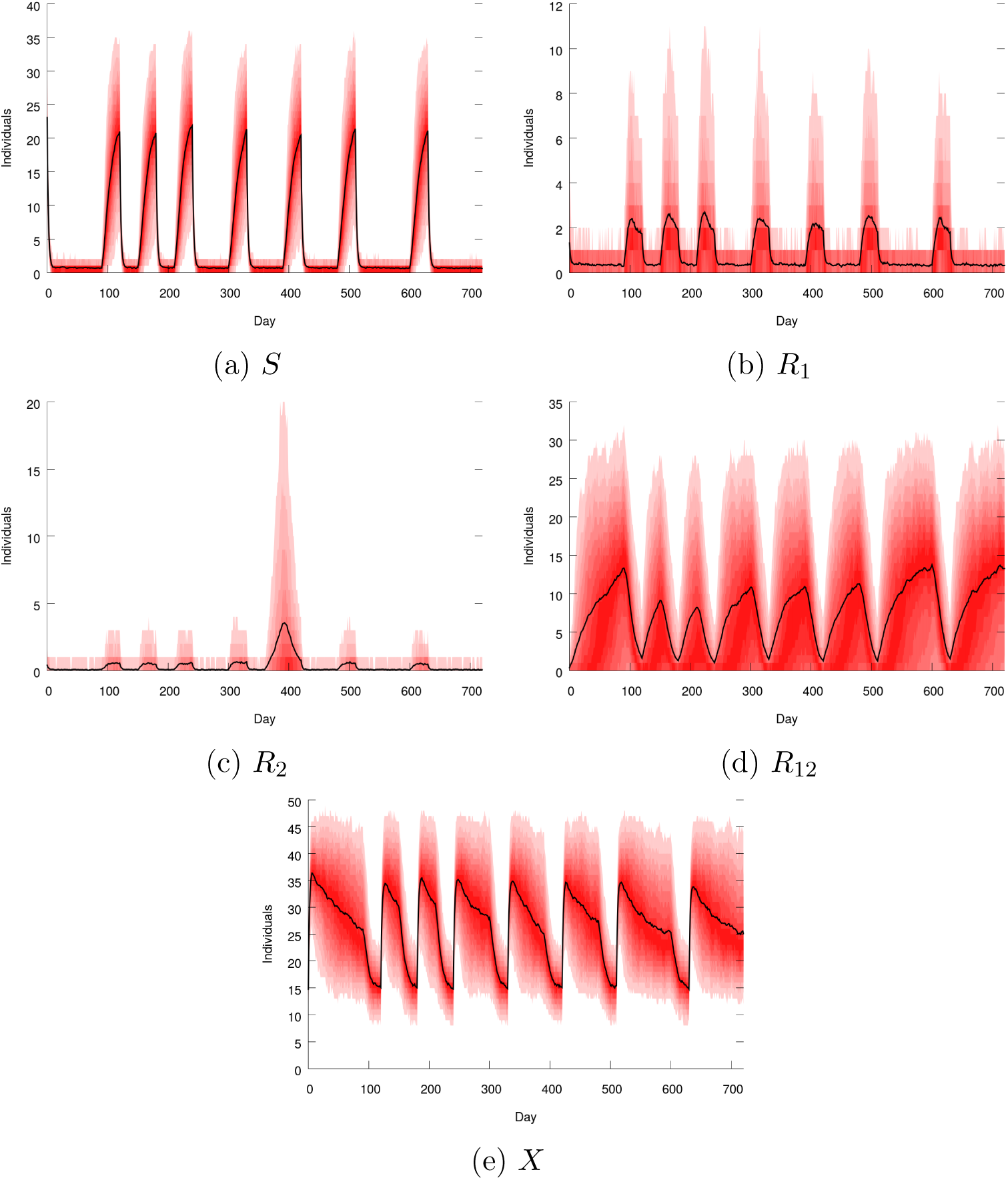
Number of patients in each compartment as a function of time when the OPTI policy is used. *Black:* average over 400 simulations. *Shades of red: i*^th^ and (100 *- i*)^th^ percentiles for *i* from 5 to 50 by increments of 5.

**Figure A.4:**
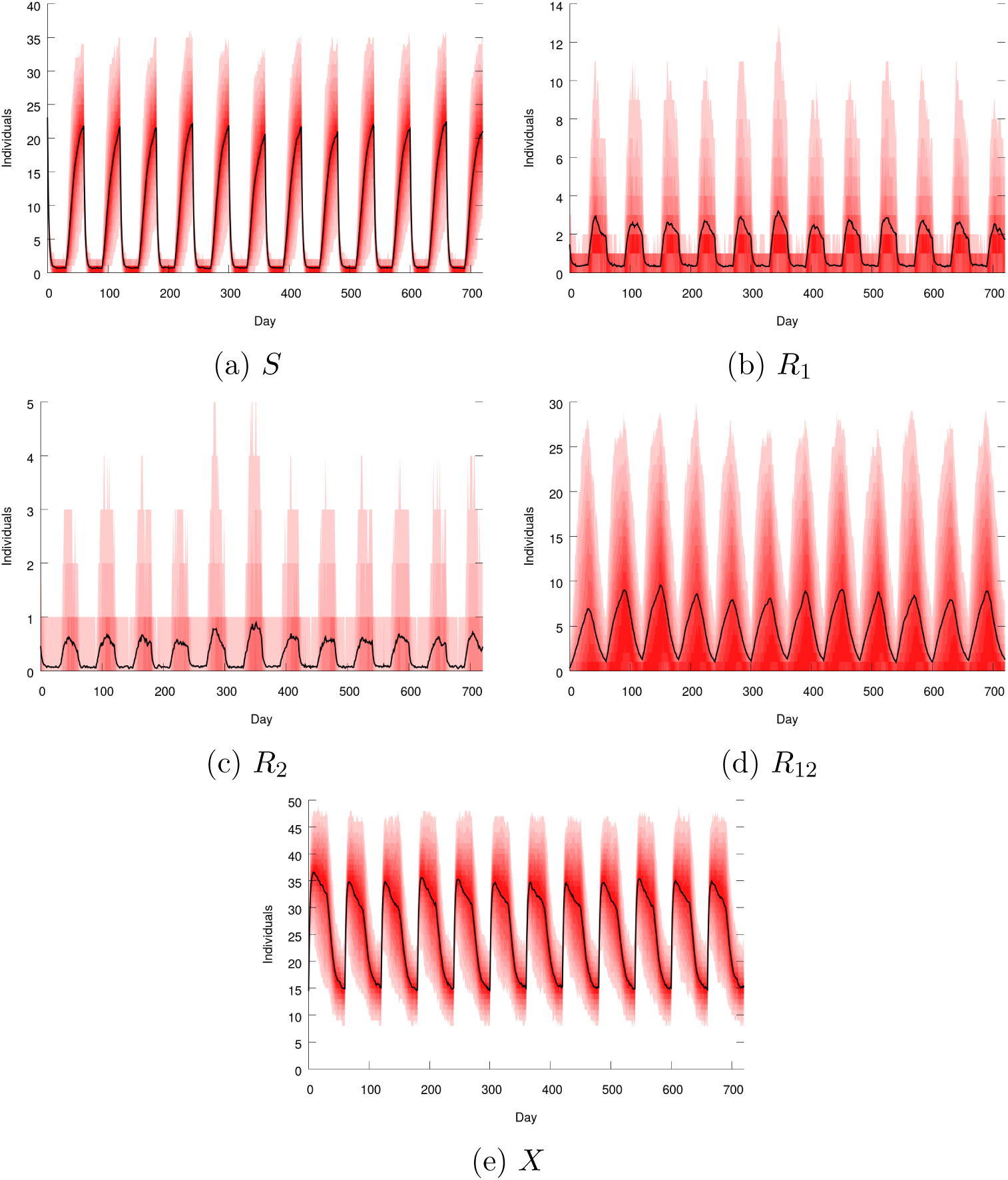
Number of patients in each compartment as a function of time when the METRO-30 policy is used. *Black:* average over 400 simulations. *Shades of red: i*^th^ and (100 *- i*)^th^ percentiles for *i* from 5 to 50 by increments of 5.

**Figure A.5:**
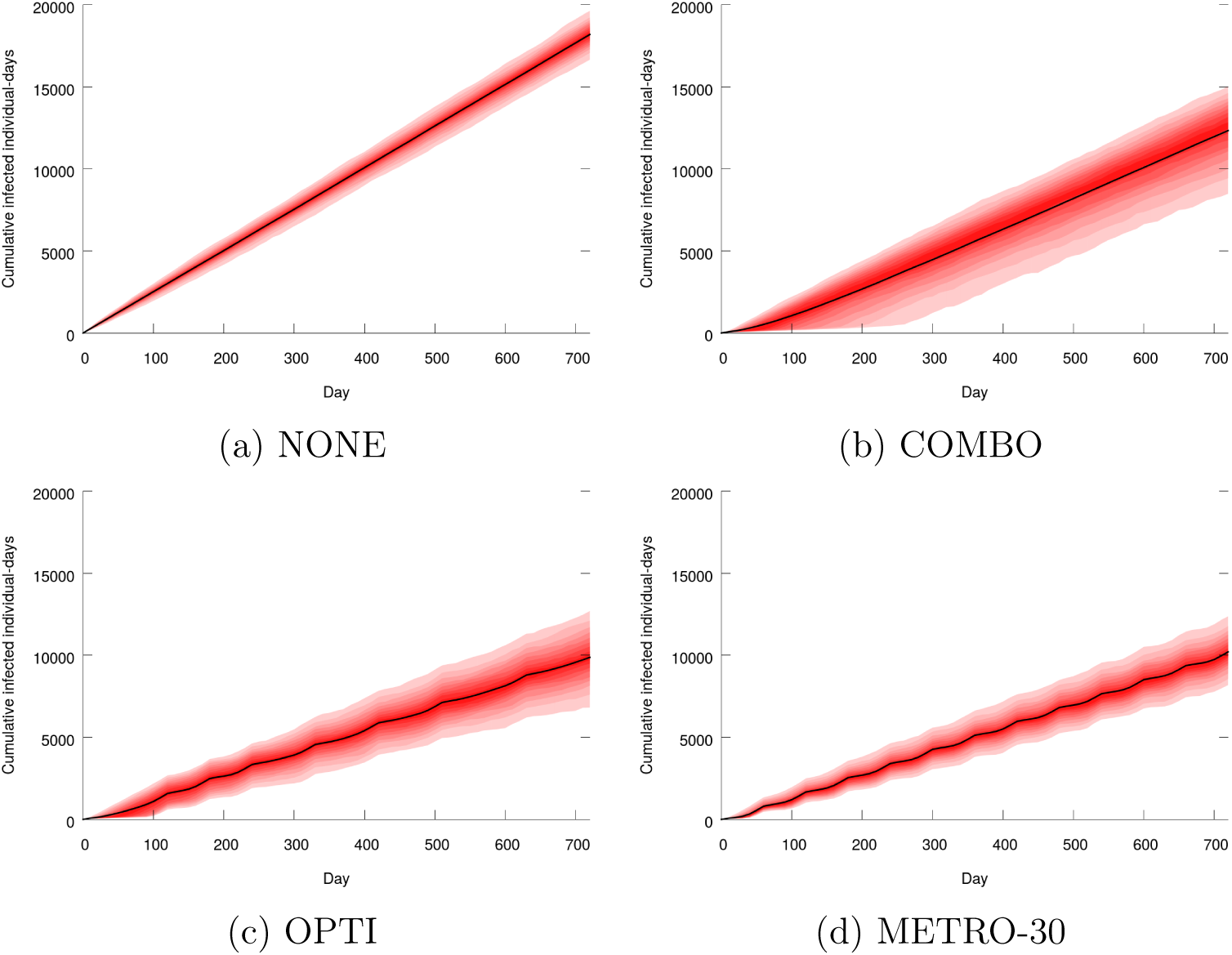
Cumulative infected patient-days as a function time when the NONE, COMBO, OPTI and METRO-30 policies are used. *Black:* average over 400 simulations. *Shades of red: i*^th^ and (100 *- i*)^th^ percentiles for *i* from 5 to 50 by increments of 5.

## Footnotes

1 For the within-host level, see [42].

2 Applications of mathematical models of health care facility infections include economic analyses [34], and complements to purely statistical approaches [43].

3 In [55], the patients benefit from the protection of their commensal microflora but this protection is removed by long-term antimicrobial treatments.

4 Notice however that our formulation disregards the respective population sizes of the new and resident strains.

5 A similar framework including a compartmental model of the community can be found in [25].

## References

[1] Dan I. Andersson. The biological cost of mutational antibiotic resistance: any practical conclusions? Current opinion in microbiology, 9(5):461–465, 2006.

[2] Dan I. Andersson and Diarmaid Hughes. Antibiotic resistance and its cost: is it possible to reverse resistance? Nature Reviews Microbiology, 8(4):260, 2010.

[3] Dan I. Andersson and Bruce R. Levin. The biological cost of antibiotic resistance. Current opinion in microbiology, 2(5):489–493, 1999.

[4] D. J. Austin, M. Kakehashi, and R. M. Anderson. The transmission dynamics of antibiotic-resistant bacteria: the relationship between resistance in commensal organisms and antibiotic consumption. Proceedings of the Royal Society of London. Series B: Biological Sciences, 264(1388):1629–1638, Nov 1997.

[5] Daren J Austin, Karl G Kristinsson, and Roy M Anderson. The relationship between the volume of antimicrobial consumption in human communities and the frequency of resistance. Proceedings of the National Academy of Sciences, 96(3):1152–1156, 1999.

[6] Miriam Barlow. What antimicrobial resistance has taught us about horizontal gene transfer. In Horizontal Gene Transfer, pages 397–411. Springer, 2009.

[7] Michael Baym, Laura K Stone, and Roy Kishony. Multidrug evolutionary strategies to reverse antibiotic resistance. Science, 351(6268):aad3292, 2016.

[8] Robert Eric Beardmore, Rafael Peña-Miller, Fabio Gori, and Jonathan Iredell. Antibiotic cycling and antibiotic mixing: which one best mitigates antibiotic resistance? Molecular biology and evolution, 34(4):802–817, 2017.

[9] Carl T Bergstrom, Monique Lo, and Marc Lipsitch. Ecological theory suggests that antimicrobial cycling will not reduce antimicrobial resistance in hospitals. Proceedings of the National Academy of Sciences, 101(36):13285–13290, 2004.

[10] Sebastian Bonhoeffer, Marc Lipsitch, and Bruce R Levin. Evaluating treatment protocols to prevent antibiotic resistance. Proceedings of the National Academy of Sciences, 94(22):12106–12111, 1997.

[11] Sebastian Bonhoeffer, Pia Abel zur Wiesch, and Roger D Kouyos. Rotating antibiotics does not minimize selection for resistance. Math Biosci Eng, 7(4):919–22, 2010.

[12] Cameron B. Browne, Edward Powley, Daniel Whitehouse, Simon M. Lucas, Peter I. Cowling, Philipp Rohlfshagen, Stephen Tavener, Diego Perez, Spyridon Samothrakis, and Simon Colton. A survey of monte carlo tree search methods. IEEE Transactions on Computational Intelligence and AI in games, 4(1):1–43, 2012.

[13] Ellsworth M Campbell and Lin Chao. A population model evaluating the consequences of the evolution of double-resistance and tradeoffs on the benefits of two-drug antibiotic treatments. PloS one, 9(1):e86971, 2014.

[14] Alessandro Cassini, Liselotte Diaz Högberg, Diamantis Plachouras, Annalisa Quattrocchi, Ana Hoxha, Gunnar Skov Simonsen, Mélanie Colomb-Cotinat, Mirjam E Kretzschmar, Brecht Devleesschauwer, Michele Cecchini, et al. Attributable deaths and disability-adjusted life-years caused by infections with antibiotic-resistant bacteria in the eu and the european economic area in 2015: a population-level modelling analysis. The Lancet Infectious Diseases, 19(1):56–66, 2019.

[15] CDC. Healthcare-associated infections. https://www.cdc.gov/hai/index.html. Accessed: 2019-03-17.

[16] BS Cooper, GF Medley, and GM Scott. Preliminary analysis of the transmission dynamics of nosocomial infections: stochastic and management effects. Journal of Hospital Infection, 43(2):131–147, 1999.

[17] Chikara Furusawa, Takaaki Horinouchi, and Tomoya Maeda. Toward prediction and control of antibiotic-resistance evolution. Current opinion in biotechnology, 54:45–49, 2018.

[18] Hajo Grundmann and B Hellriegel. Mathematical modelling: a tool for hospital infection control. The Lancet infectious diseases, 6(1):39–45, 2006.

[19] Erik Gullberg, Sha Cao, Otto G. Berg, Carolina Ilbäck, Linus Sandegren, Diarmaid Hughes, and Dan I. Andersson. Selection of resistant bacteria at very low antibiotic concentrations. PLOS Pathogens, 7(7):1–9, 07 2011.

[20] Nicolas Houy and François Le Grand. Optimizing immune cell therapies with artificial intelligence. Journal of Theoretical Biology, 461:34–40, jan 2019.

[21] Nicolas Houy and François Le Grand. Optimal dynamic regimens with artificial intelligence: The case of temozolomide. PLOS ONE, 13(6):1–15, 06 2018.

[22] Pentti Huovinen. Mathematical model—tell us the future! Journal of Antimicrobial Chemotherapy, 56(2):257–258, 2005.

[23] Annette Jepson. Microbiology and infection control. In Carlos M H Gómez, editor, Clinical Intensive Care Medicine, chapter 10. Imperial College Press, 2014.

[24] Roger D. Kouyos, Pia Abel zur Wiesch, and Sebastian Bonhoeffer. Informed switching strongly decreases the prevalence of antibiotic resistance in hospital wards. PLoS computational biology, 7(3):e1001094, 2011.

[25] Roger D. Kouyos, Pia Abel zur Wiesch, and Sebastian Bonhoeffer. On being the right size: the impact of population size and stochastic effects on the evolution of drug resistance in hospitals and the community. PLoS pathogens, 7(4):e1001334, 2011.

[26] BR Levin, M Lipsitch, V Perrot, S Schrag, R Antia, Lone Simonsen, N Moore Walker, and FM Stewart. The population genetics of antibiotic resistance. Clinical infectious diseases, 24(Supplement_1):S9–S16, 1997.

[27] Bruce R Levin and Marc JM Bonten. Cycling antibiotics may not be good for your health. Proceedings of the National Academy of Sciences, 101(36):13101–13102, 2004.

[28] Marc Lipsitch and Carl T Bergstrom. Modeling of antibiotic resistance in the icuus slant. In R. A. Weinstein and M. Bonten, editors, Infection Control in the ICU environment. Kluwer, 2002.

[29] Marc Lipsitch, Carl T Bergstrom, and Bruce R Levin. The epidemiology of antibiotic resistance in hospitals: paradoxes and prescriptions. Proceedings of the National Academy of Sciences, 97(4):1938–1943, 2000.

[30] Marc Lipsitch and Matthew H. Samore. Antimicrobial use and antimicrobial resistance: a population perspective. Emerging infectious diseases, 8(4):347–354, 2002.

[31] Eduardo Massad, Sigfrid Lundberg, and Hyun Mo Yang. Modeling and simulating the evolution of resistance against antibiotics. International journal of bio-medical computing, 33(1):65–81, 1993.

[32] Anita H. Melnyk, Alex Wong, and Rees Kassen. The fitness costs of antibiotic resistance mutations. Evolutionary applications, 8(3):273–283, 2015.

[33] P. A. Milligan, M. J. Brown, B. Marchant, S. W. Martin, P. H. van der Graaf, N. Benson, G. Nucci, D.J. Nichols, R. A. Boyd, J. W. Mandema, S. Krishnaswami, S. Zwillich, D. Gruben, R. J. Anziano, T. C. Stock, and R. L. Lalonde. Model-based drug development: A rational approach to efficiently accelerate drug development. Clinical Pharmacology & Therapeutics, 93(6):502–514, mar 2013.

[34] Richard E. Nelson, Rishi Deka, Karim Khader, Vanessa W. Stevens, Marin L. Schweizer, and Michael A. Rubin. Dynamic transmission models for economic analysis applied to health care-associated infections: A review of the literature. American Journal of Infection Control, 45(12):1382–1387, 2017.

[35] Lindsay E. Nicolle. Infection control programmes to contain antimicrobial resistance. WHO, 2001. Available at https://www.who.int/csr/resources/publications/drugresist/infection_control.pdf.

[36] Michael S. Niederman. Is “crop rotation” of antibiotics the solution to a “resistant” problem in the icu? American Journal of Respiratory and Critical Care Medicine, 156(4):1029–1031, 1997.

[37] Uri Obolski and Lilach Hadany. Implications of stress-induced genetic variation for minimizing multidrug resistance in bacteria. BMC medicine, 10(1):89, 2012.

[38] Uri Obolski, Gideon Y Stein, and Lilach Hadany. Antibiotic restriction might facilitate the emergence of multi-drug resistance. PLoS computational biology, 11(6):e1004340, 2015.

[39] Zaheerabbas Patwa and Lindi M. Wahl. The fixation probability of beneficial mutations. Journal of The Royal Society Interface, 5(28):1279–1289, 2008.

[40] R Peña-Miller and Robert Beardmore. Rotating antibiotics selects optimally against antibiotic resistance, in theory. Mathematical Biosciences & Engineering, 7(3):pp. 527–552, 2010.

[41] Rafael Peña-Miller and Robert Beardmore. Antibiotic cycling versus mixing: the difficulty of using mathematical models to definitively quantify their relative merits. Mathematical Biosciences & Engineering, 7(4):923–933, 2010.

[42] Rafael Peña-Miller, David Lähnemann, Hinrich Schulenburg, Martin Ackermann, and Robert Beardmore. The optimal deployment of synergistic antibiotics: a control-theoretic approach. Journal of The Royal Society Interface, 9(75):2488–2502, 2012.

[43] Rebecca A Pierce, Justin Lessler, and Aaron M Milstone. Expanding the statistical toolbox: analytic approaches for cohort studies with healthcare-associated infectious outcomes. Current opinion in infectious diseases, 28(4):384, 2015.

[44] D. E. Ramsay, J. Invik, S. L. Checkley, S. P. Gow, N. D. Osgood, and C. L. Waldner. Application of dynamic modelling techniques to the problem of antibacterial use and resistance: a scoping review. Epidemiology & Infection, 146(16):2014–2027, 2018.

[45] Timothy C. Reluga. Simple models of antibiotic cycling. Mathematical medicine and biology: a journal of the IMA, 22(2):187–208, 2005.

[46] Damien Roux, Olga Danilchanka, Thomas Guillard, Vincent Cattoir, Hugues Aschard, Yang Fu, Francois Angoulvant, Jonathan Messika, Jean-Damien Ricard, John J Mekalanos, et al. Fitness cost of antibiotic susceptibility during bacterial infection. Science translational medicine, 7(297):297ra114–297ra114, 2015.

[47] Anna Seigal, Portia Mira, Bernd Sturmfels, and Miriam Barlow. Does antibiotic resistance evolve in hospitals? Bulletin of mathematical biology, 79(1):191–208, 2017.

[48] Ian H Spicknall, Betsy Foxman, Carl F Marrs, and Joseph NS Eisenberg. A modeling framework for the evolution and spread of antibiotic resistance: literature review and model categorization. American journal of epidemiology, 178(4):508–520, 2013.

[49] Evelina Tacconelli and Maria Diletta Pezzani. Public health burden of antimicrobial resistance in europe. The Lancet Infectious Diseases, 19(1):4–6, 2019.

[50] Fred C Tenover. Mechanisms of antimicrobial resistance in bacteria. The American journal of medicine, 119(6):S3–S10, 2006.

[51] Burcu Tepekule, Hildegard Uecker, Isabel Derungs, Antoine Frenoy, and Sebastian Bonhoeffer. Modeling antibiotic treatment in hospitals: A systematic approach shows benefits of combination therapy over cycling, mixing, and mono-drug therapies. PLoS computational biology, 13(9):e1005745, 2017.

[52] Hildegard Uecker and Sebastian Bonhoeffer. Modeling antimicrobial cycling and mixing: Differences arising from an individual-based versus a population-based perspective. Mathematical biosciences, 294:85–91, 2017.

[53] Esther van Kleef, Julie V Robotham, Mark Jit, Sarah R Deeny, and William J Edmunds. Modelling the transmission of healthcare associated infections: a systematic review. BMC infectious diseases, 13(1):294, 2013.

[54] Christian JH von Wintersdorff, John Penders, Julius M van Niekerk, Nathan D Mills, Snehali Majumder, Lieke B van Alphen, Paul HM Savelkoul, and Petra FG Wolffs. Dissemination of antimicrobial resistance in microbial ecosystems through horizontal gene transfer. Frontiers in microbiology, 7:173, 2016.

[55] Pia Abel zur Wiesch, Roger Kouyos, Sören Abel, Wolfgang Viechtbauer, and Sebastian Bonhoeffer. Cycling empirical antibiotic therapy in hospitals: meta-analysis and models. PLoS pathogens, 10(6):e1004225, 2014.

[56] Pia Abel zur Wiesch, Roger Kouyos, Jan Engelstädter, Roland R Regoes, and Sebastian Bonhoeffer. Population biological principles of drug-resistance evolution in infectious diseases. The Lancet infectious diseases, 11(3):236–247, 2011.

